# Neuropilin-1 functions as a proviral and immunoregulatory host factor during Chikungunya virus infection

**DOI:** 10.64898/2026.07.27.741004

**Authors:** Kshyama Subhadarsini Tung, Chandan Mahish, Soumyajit Ghosh, Kaustuv Mukherjee, Sharad Singh, Bijita Bhowmick, Somlata Khamaru, Harshit Borasi, Mahendra Gaur, Bharat Bhusan Subudhi, Soma Chattopadhyay, Subhasis Chattopadhyay

**Author notes:** **Correspondence: Dr. Subhasis Chattopadhyay** School of Biological Sciences National Institute of Science Education and Research Bhubaneswar Jatni, khorda, Odisha, India- 752050. Present address: National Institute of Allergy and Infectious Diseases, NIH, USA. Present address: Department of Zoology, Burdwan University, West Bengal, India (Somlata Khamaru). Equal Contribution (Chandan Mahish, Soumyajit Ghosh).

## Abstract

Neuropilin-1 (NRP1) is a transmembrane glycoprotein involved in angiogenesis, neurodevelopment, inflammation, cancer driven immune suppression, and immune homeostasis. However, its contribution to virus-induced immune responses is not explored. Chikungunya virus (CHIKV) is a re-emerging arthritogenic alphavirus that causes severe arthralgia, and myalgia, accompanied by heightened inflammatory cytokine responses. The host factors that drive these inflammatory responses, however, remain poorly defined. Here, in this current study, the role of NRP1 in CHIKV infection was investigated using *in vitro*, and *in vivo* model systems. Using genetic manipulation, pharmacological, and antibody blockade-mediated approaches in murine and human cellular infection models, it was demonstrated that NRP1 promotes CHIKV infection while restraining the production of proinflammatory cytokines. Furthermore, NRP1 inhibition selectively increased JNK phosphorylation. Hence, inhibiting JNK reduced the elevated cytokine production caused by NRP1 blockade. Moreover, NRP1 interacted with CHIKV-E1 and is involved in multiple phases of CHIKV infection. In addition, it was demonstrated that NRP1 inhibition using EG00229 trifluoroacetate can reduce viral infection in several CHIKV-susceptible cells, human peripheral blood macrophages, and in the *in vivo* mice model of infection. Together, these findings indicate that NRP1 is an important host factor and a probable therapeutic target during CHIKV infection.

**IMPORTANCE:** Chikungunya virus (CHIKV) causes acute febrile illness that can progress to debilitating chronic musculoskeletal and occasional neurological complications. The absence of a globally available effective vaccine and the lack of specific antivirals underscore CHIKV as a major burden, especially in endemic tropical regions. There is a growing need to understand host factors that shape CHIKV-driven immune response. The importance of our study lies in understanding the immunoregulatory role of the host receptor Neuropilin-1 (NRP1) during CHIKV infection. We identify NRP1 as a novel host factor of CHIKV infection that also acts as a rheostat to restrict the inflammatory viral immune response. Moreover, we report anti-viral potential of the NRP1 antagonist EG00229 trifluoroacetate in multiple cell lines, primary cells, and mice model. Our findings indicate NRP1 as a probable therapeutic target in CHIKV pathogenesis.

## 1. INTRODUCTION

Chikungunya virus (CHIKV) is a mosquito-borne alphavirus of the togaviridae family. Since its global re-emergence, CHIKV has caused prevalent outbreaks across the world and imposes a significant public health burden (1). In spite of advances in CHIKV vaccine development, including VLA1553 (Ixchiq, live-attenuated), and virus-like particle chikungunya vaccine (VIMKUNYA), no universally deployable solutions yet exist (2). Moreover, the lack of specific antivirals underscores the critical need to study host factors that regulate CHIKV pathogenesis. CHIKV causes acute polyarthralgia and myalgia with long-term musculoskeletal complications and exhibits broad cellular tropism, infecting epithelial, endothelial, muscle, fibroblast, and immune cells (3). CHIKV infection leads to production of robust proinflammatory cytokines (e.g. TNF, IL-6, etc.) (4, 5). CHIKV disease severity depends on the balance between antiviral responses and pathological inflammation (6). The early antiviral immune activation correlates with viral control; however excessive or prolonged inflammation drives chronic disease (7, 8). Immune cells, primarily monocytes and macrophages, initiate antiviral responses via pattern recognition receptors (PRRs), producing inflammatory cytokines and chemokines (9). While, monocytes and macrophages mediate anti-viral defenses, they also support viral persistence, particularly in joints, contributing to long term complications (8, 10). In addition, depletion of inflammatory macrophage subsets (CD64⁺MHCII⁺) reduces joint pathology, while specific monocyte populations (Ly6C^hi^CCR2**^+^**) drive protective interferon responses during CHIKV infection (11, 12). Furthermore, elevated inflammatory markers in patient peripheral blood mononuclear cells (PBMCs) correlate with disease severity, underscoring the central role of monocytes/macrophages in CHIKV immunopathogenesis (13). CHIKV infection is characterized by dysregulated innate and adaptive immune responses; however, host factors that coordinate viral infection and immune responses still remain largely unexplored.

Neuropilin-1 (NRP1) is a multifunctional transmembrane glycoprotein originally identified as a receptor for class III semaphorins and vascular endothelial growth factors, with established roles in neuronal guidance and angiogenesis (14). NRP1 coordinates dendritic cell-T-cell interactions, regulatory T-cell stability, macrophage migration, and shapes immune responses and tolerance (14). NRP1 depletion or inhibition enhances pro-inflammatory signaling, cytokine production, and inflammasome activation across multiple disease models (15–21). Myeloid specific knockout of NRP1 leads to hyperinflammation, exacerbated sepsis, and increased IL-6, IL-1β, and TNF-α production via TLR4-NF-κB pathways (15). Furthermore, targeting NRP1⁺ Tregs in the tumor microenvironment reduces immunosuppression and improves antitumor immunity (22). Collectively, NRP1 appears to be a critical immunoregulatory molecule, with its dysregulated expression consistently associated with loss of immune regulation across diseases (15, 16, 23, 24).

Beyond its canonical role, NRP1 has currently emerged as a host entry factor utilized by different viruses. It has been implicated in facilitating viral entry and attachment of Epstein-Barr virus (EBV), SARS-CoV-2, human T-cell leukemia virus type-1 (HTLV-1), cytomegalovirus, hepatitis B virus (HBV), Kaposi’s sarcoma-associated herpesvirus (KSHV), and pseudorabies virus (25–31). However, its involvement in alphavirus infection, including CHIKV, remains undefined. In addition, its role in regulating host immune responses during any viral infection is yet to be explored. Given its established functions in immune regulation and disease pathogenesis, the potential involvement of NRP1 in CHIKV-induced immune responses requires critical investigation.

In this study, the role of NRP1 in CHIKV immunopathogenesis was investigated using *in vitro* and *in vivo* models of infection. Accordingly, it was demonstrated that NRP1 contributes to CHIKV infection while also regulating the host inflammatory responses.

## 2. RESULTS

### 2.1. NRP1 expression is downregulated during CHIKV infection in macrophages, *in vitro*

To investigate whether CHIKV infection alters NRP1 expression, surface and total NRP1 levels were assessed in RAW 264.7 macrophages following infection. RAW 264.7 cells were infected with CHIKV at MOI 5 and harvested at 8hpi as mentioned previously (5, 32), followed by flow cytometry-based analysis of surface NRP1 expression. It was observed that CHIKV infection led to a significant downregulation of NRP1 surface expression (**Fig. S1A**). Consistent with these observations, Western blot analysis of cell lysates also revealed a reduction in total NRP1 protein levels in infected macrophages (**Fig. S1B**). Collectively, these findings indicate that CHIKV infected macrophages show reduced NRP1 expression.

### 2.2. Functional inhibition of NRP1 restricts CHIKV infection in macrophages

To investigate the functional role of NRP1 during CHIKV infection, NRP1 was pharmacologically inhibited using EG00229 trifluoroacetate (EG). An MTT assay for EG was conducted on RAW 264.7 and PMA-differentiated THP-1 cells (dTHP-1) to assess possible cytotoxicity. Both cell types were found to be more than 90% viable up to 10µM concentration of EG **(Fig. S1C-D)** and this dose was selected for further study. EG was administered before, during, or after CHIKV infection. Cells were analyzed by flow cytometry to assess viral infection, while culture supernatants were subjected to plaque assay to quantify viral titers. Viral infection was assessed via flow cytometry by using either the envelope protein E2, or non-structural protein nsP2 of CHIKV as reliable markers (33, 34). Additionally, whole cell lysates were analyzed by Western blot to assess the levels of CHIKV protein E2. Treatment of RAW 264.7 cells with 10µM of EG treatment resulted in a reduction of infection in terms of % positive cells for CHIKV E2 (19.9% to 15%) and CHIKV nsP2 (15.9% to 11.4%) **(Fig. 1A-D)**. To further validate the effects of NRP1 inhibition on CHIKV infection, membrane NRP1 was also blocked with an anti-NRP1 antibody prior to CHIKV infection. A decrease in the percent positive cells for CHIKV E2 (19% to 11.9%) were observed in cells treated with anti-NRP1 antibody as compared to the control IgG antibody (**Fig. 1E-G**). Furthermore, a reduction in viral titer was observed due to EG treatment (∼2.51-fold reduction, approximately 60.2%) (**Fig. 1H**), and anti-NRP1 blockade (∼2.56-fold reduction, approximately 61.0%) in CHIKV infected RAW 264.7 cells (**Fig. 1I**). Western blot analysis showed a reduction in CHIKV E2 protein levels in the presence of EG treatment and anti-NRP1 antibody blockade **(Fig. 1J)**. EG treatment did not have a significant effect on surface NRP1 expression in RAW 264.7 cells, as assessed by flow cytometry **(Fig. S1I)**. However, EG treatment resulted in a modest, albeit non-significant, increase in surface NRP1 expression compared with CHIKV-infected macrophages (**Fig. S1J**). In contrast, Western blot analysis demonstrated a modest but significant increase in total NRP1 protein levels in CHIKV-infected RAW 264.7 macrophages following EG treatment. (**Fig. S1K**).

**Figure 1:**
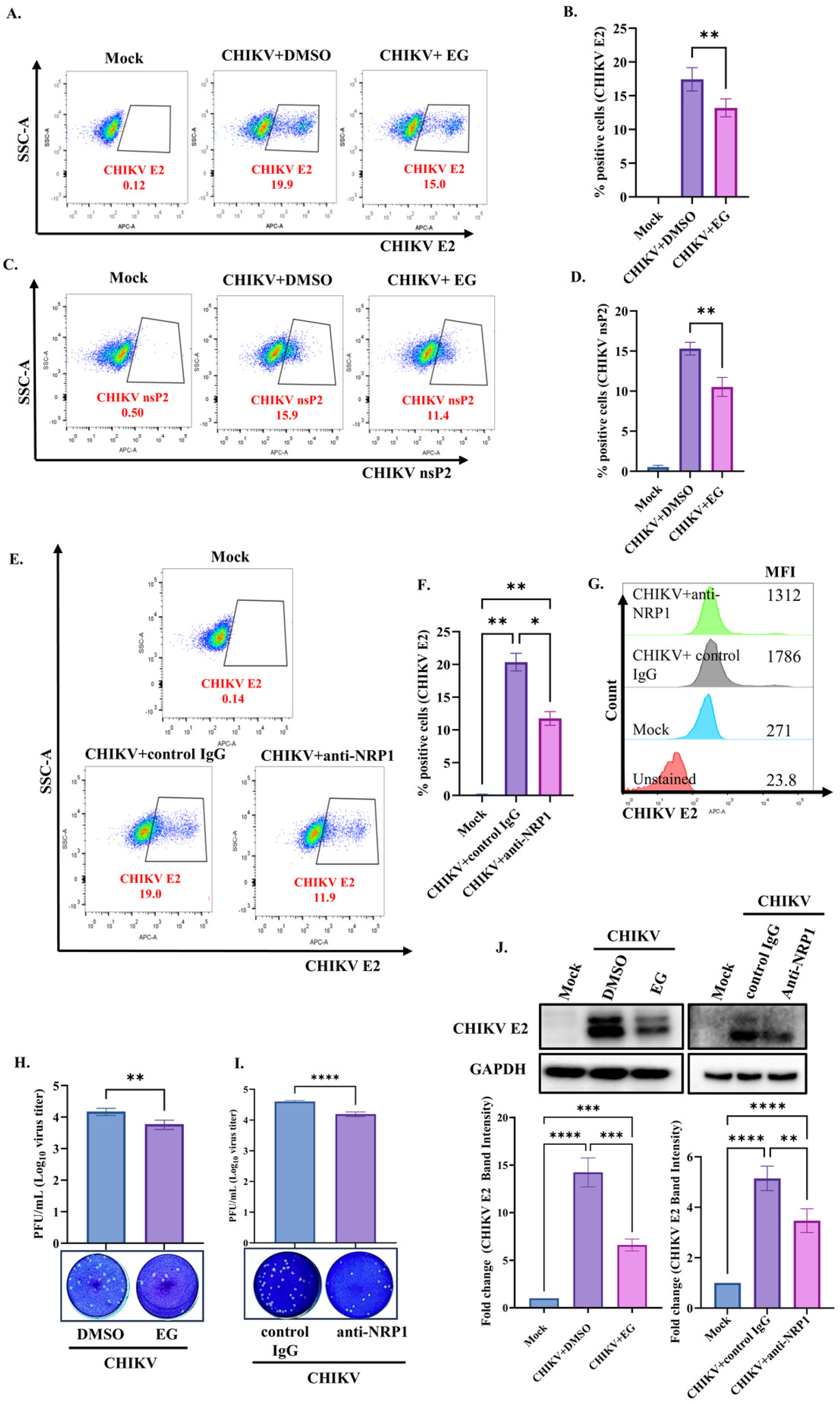
Pharmacological inhibition or antibody blockade of NRP1 reduces CHIKV infection in macrophages *in vitro*. RAW 264.7 cells were treated with DMSO or EG00229 trifluoroacetate (EG). For antibody-mediated blockade, cells were treated with anti-NRP1 or isotype control IgG antibodies. Cells were infected with CHIKV at an infection multiplicity (MOI) of 5 for 2 h and harvested at 8 h post-infection (hpi). **(A)** Flow cytometry dot plots, and (**B)** representative bar graph showing the percentage of CHIKV E2-positive cells **(C)** Flow cytometry dot plots, and **(D)** representative bar graph showing the percentage of CHIKV nsP2-positive cells. **(E)** Flow cytometry dot plots, and **(F)** representative bar graph showing the percentage of CHIKV E2-positive cells. **(G)** Mean fluorescence intensity (MFI) of CHIKV E2 following NRP1 inhibition by antibody blockade. Bar graph representing CHIKV titers following **(H)** EG treatment and **(I)** anti-NRP1 antibody blockade, as determined by plaque assay. **(J)** Representative western blots and bar graphs (densitometry of E2 band intensity) showing CHIKV E2 protein levels following NRP1 inhibition by EG or anti-NRP1 antibody. Data are presented as the mean ± SD from three independent experiments. Statistical significance was determined as indicated (ns, not significant; *, P < 0.05; **, P ≤ 0.01; ***, P ≤ 0.001; ***, P ≤ 0.0001).

To extend these findings in human macrophages; the effect of NRP1 inhibition was additionally examined in dTHP-1 cells. NRP1 inhibition led to a ∼65% reduction of CHIKV infection in dTHP-1 cells as evident by decreased % positive cells for CHIKV E2 (32.6% to 11.2%) and CHIKV nsP2 (27% to 8.76%) (**Fig. S2A-D**). Taken together, these results suggest that NRP1 inhibition reduces CHIKV infection in macrophages.

### 2.3. NRP1 promotes CHIKV infection across diverse cell types

To assess whether NRP1 promotes CHIKV infection in multiple cell types, CHIKV infection was examined following EG treatment in BV2 microglial, Vero epithelial, and C2C12 muscle cells. The non-cytotoxic concentration of EG was determined in all cell lines using the MTT assay (**Fig. S1E-G**). Accordingly, 10µM EG was selected for subsequent experiments. EG was administered before, during, as well as in post-infection phase up to appropriate time points. Accordingly, C2C12 (MOI 0.05), BV2 (MOI 5), and Vero cells (MOI 0.1) were infected and harvested at 15hpi, 8hpi, and 12hpi respectively. CHIKV infected and EG treated cells were subjected to flow cytometry to assess CHIKV infection. EG directed NRP1 inhibition led to significant reduction in percentage of CHIKV nsP2 positive cells (48.7% to 31.4%) in murine muscle cells (**Fig. 2A-B**), and decreased viral titer (∼2.31-fold reduction, approximately 56.8% decrease) (**Fig. 2C**). Similarly, EG treatment reduced % CHIKV nsP2 positive cells (15.6% to 9.95%) and decreased viral titer in BV2 cells (∼1.87-fold reduction, approximately 46.4% decrease) (**Fig. 2D-F**). Moreover, NRP1 inhibition also reduced % CHIKV nsP2 positive cells (27% to 19.4%), and viral titers in Vero cells (∼2.88-fold reduction, approximately 65.2% decrease) **(Fig. 2G-I).** Collectively, these observations suggest that NRP1 contributes to CHIKV infection across different cell types.

**Figure 2:**
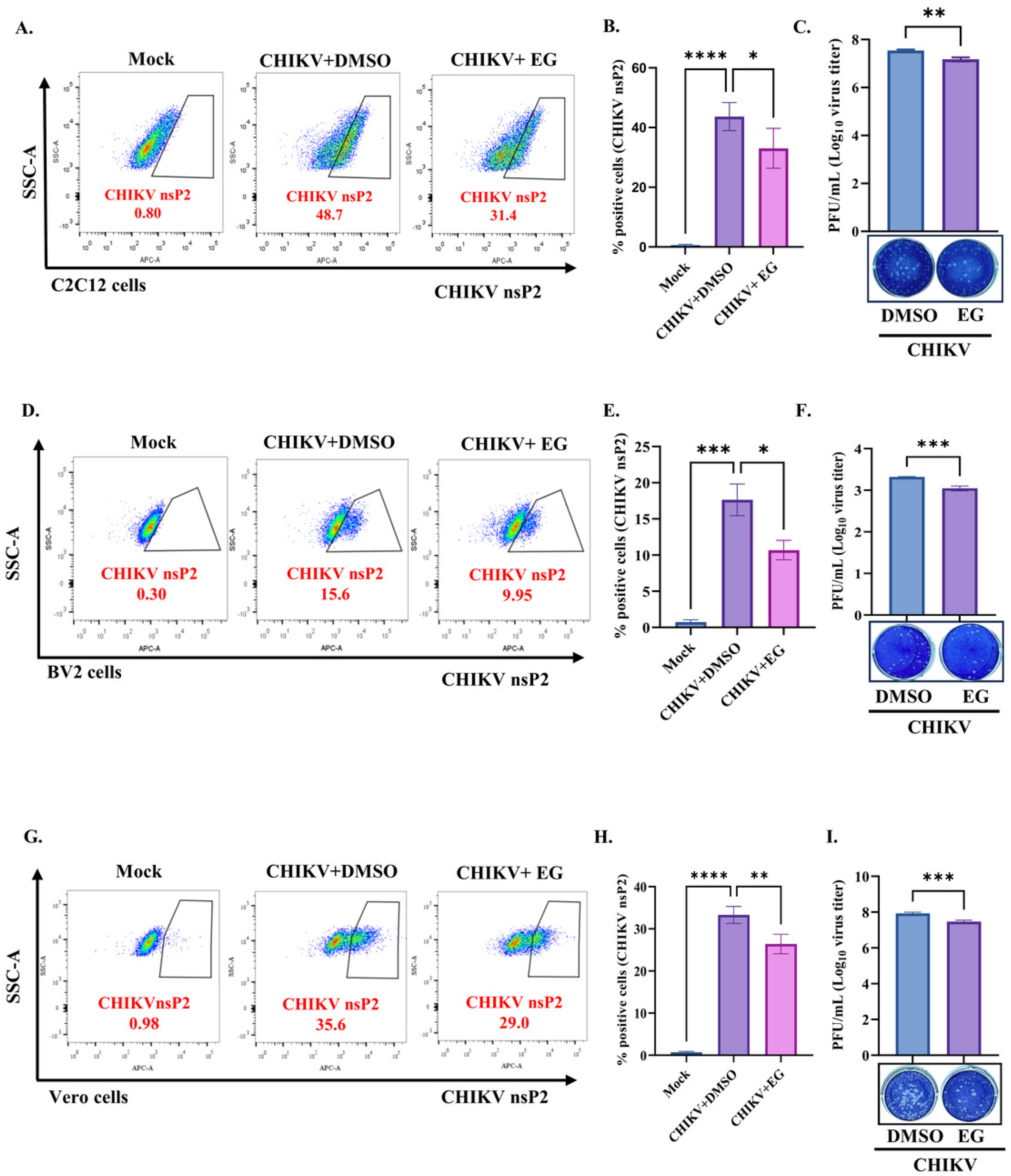
EG-directed inhibition of NRP1 reduces CHIKV infection and viral titers in C2C12, BV2, and Vero cells. CHIKV infected and EG treated cells were harvested and subjected to flow cytometry to assess infection **(A)** Flowcytometric dot plots, and **(B)** representative bar graph showing percentage of positive cells for CHIKV nsP2, and **(C)** Bar graph depicting viral titer in C2C12 cells with NRP1 inhibition. **(D)** Flowcytometric dot plots, and **(E)** representative bar graph of % positive cells for CHIKV nsP2, and **(F)** Bar graph depicting viral titer in BV2 cells with EG treatment. **(G)** Flowcytometric dot plots, and **(H)** representative bar graph of percentage of positive cells for CHIKV nsP2, and **(I)** Bar graph depicting viral titer in Vero cells with EG treatment. Data are presented as the mean ± SD from three independent experiments. Statistical significance was determined as indicated (ns, not significant; *, P < 0.05; **, P ≤ 0.01; ***, P ≤ 0.001; ***, P ≤ 0.0001).

### 2.4. Expression of NRP1 positively regulates CHIKV infection: knockdown restricts, while overexpression enhances infection

To assess the functional role of NRP1 in CHIKV infection, siRNA-mediated knockdown was carried out in HEK293T cells. HEK cells were transfected with 90pm of siRNA specific to NRP1 or scramble. No significant cytotoxic effects were detected in siRNA-treated cells. Significant depletion of NRP1 (∼52.3% knockdown efficiency) was confirmed by western blot analysis **(Fig. 3A-B)**. As previously observed, NRP1 expression was further reduced during CHIKV infection in knockdown cells compared with scrambled controls (**Fig. 3C**). A significant reduction in CHIKV E2 protein expression (∼39% reduction) was observed in NRP1-silenced cells compared to scrambled controls **(Fig. 3D)**. In addition, lower infectious titers (∼3.08-fold reduction, approximately 67.5% decrease) were detected upon NRP1 knockdown **(Fig. 3E).** Similar experiments were performed with murine C2C12 cells, where NRP1 knockdown was associated with reduced CHIKV E2 levels (**Fig. S2E-G)**.

**Figure 3:**
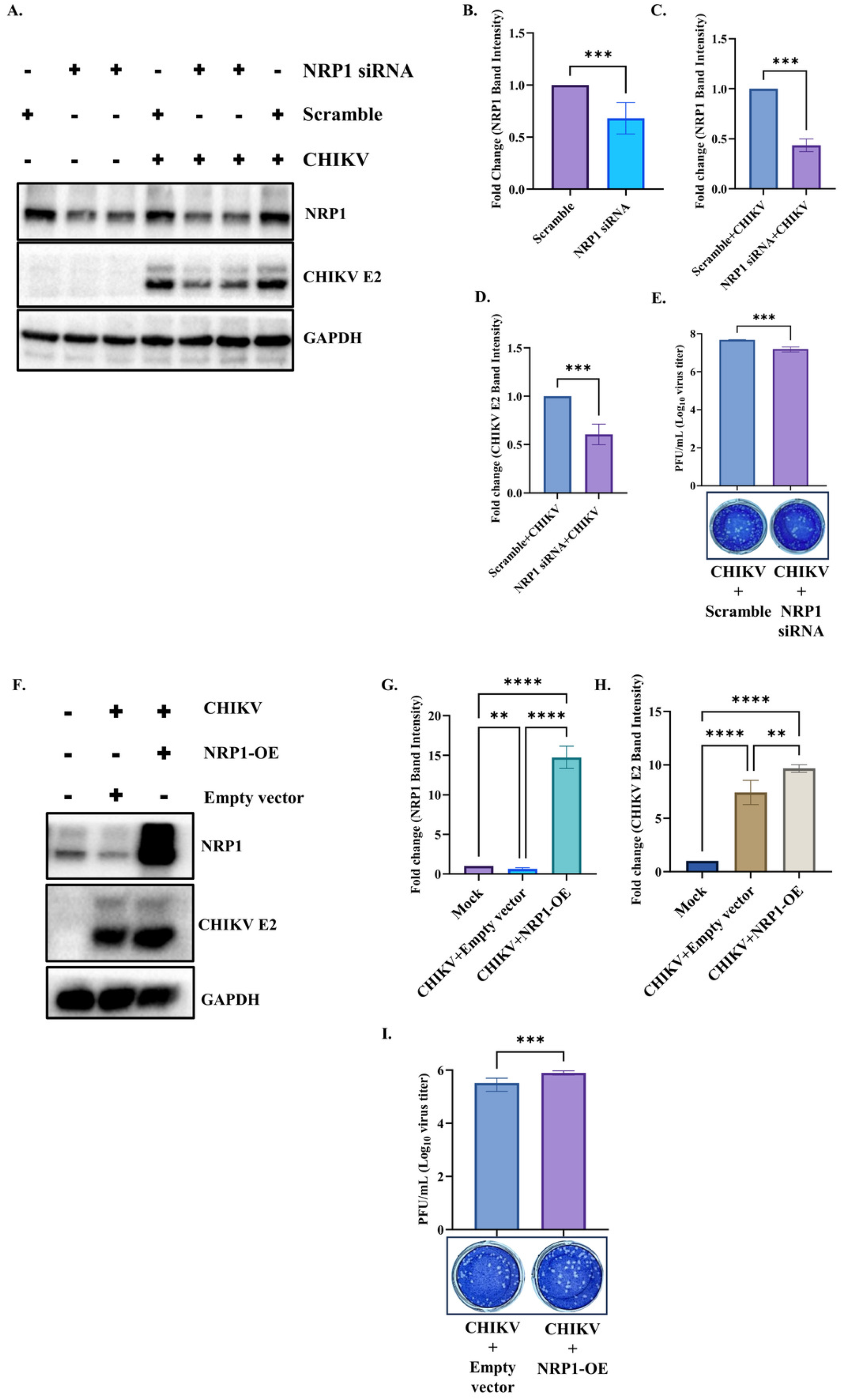
Knockdown of NRP1 restricts viral infection, and its overexpression enhances infection. HEK cells were transfected with a scramble control or NRP1-specific siRNA and then infected with CHIKV at an MOI of 0.1. Cells were harvested at 15 h post-infection (hpi) to evaluate NRP1 knockdown and its effect on viral infection. For overexpression studies, HEK cells were transfected with an empty vector or NRP1-expressing pcDNA3.1, infected with CHIKV at an MOI of 0.1, and harvested at 15 hpi. NRP1 overexpression was confirmed by immunoblotting, and its impact on CHIKV infection was assessed by Western blot and plaque assay. **(A)** Immunoblot analysis of NRP1 and CHIKV E2 protein levels in NRP1-silenced HEK cells following CHIKV infection. **(B)** Quantification of NRP1 knockdown efficiency in HEK cells. **(C)** Quantification of NRP1 levels in CHIKV infected scramble vs NRP1 knocked down cells **(D)** Densitometric analysis showing CHIKV E2 levels, and **E)** Bar graph showing viral titers in NRP1-knockdown HEK cells. **(F)** Immunoblot analysis of NRP1 and CHIKV E2 protein levels in CHIKV-infected HEK cells following NRP1 overexpression. **(G)** Fold change in NRP1 protein levels following overexpression in HEK cells (NRP1-OE), as determined by Western blotting. **(H)** Quantification of CHIKV infection in HEK cells upon NRP1 overexpression (NRP1-OE). **(I)** Bar graph showing viral titers in CHIKV-infected HEK cells with NRP1 overexpression (NRP1-OE). Data are presented as the mean ± SD from three independent experiments. Statistical significance was determined as indicated (ns, not significant; *, P < 0.05; **, P ≤ 0.01; ***, P ≤ 0.001; ***, P ≤ 0.0001).

To validate the effects of NRP1 overexpression on CHIKV infection, HEK cells were transfected with pcDNA3.1-NRP1 or an empty vector before CHIKV infection. No significant cytotoxic effects were detected in the transfected cells. NRP1 overexpression was validated by Western blot and densitometric quantification **(Fig. 3F-G)**. An increase of CHIKV E2 protein levels and higher infectious CHIKV titers (∼2.45-fold increase) were observed in NRP1 overexpressed HEK cells relative to empty vector-transfected controls **(Fig. 3H-I)**. Taken together, it can be suggested that NRP1 positively regulates CHIKV infection.

### 2.5. The NRP1 inhibitor decreases viral load by affecting different phases of CHIKV infection

To determine whether NRP1 inhibition selectively affects any specific stages of viral infection, RAW 264.7 cells were treated with EG at distinct phases of CHIKV infection: pre-treatment (2 h prior to infection), during infection, post-infection (up to 8hpi), and a combined regimen (pre + during + post). EG treatment resulted in a reduction in the percentage of CHIKV E2–positive cells across all conditions -pre, -during, -post, and, -combined as compared to the DMSO control (**Fig. 4A**). Consistently, NRP1 inhibition also decreased viral titer by 33.7%, 40%, 34.5%, and 59% in the pre-, during-, post-, and combined treatment groups, respectively (**Fig. 4B)**. Similar trends were observed in physiologically relevant CHIKV-infected C2C12 cells, where EG treatment reduced the percentage of CHIKV nsP2-positive cells across all phases of infection (**Fig. 4C**). Since EG showed anti-CHIKV activity in the “during” phase of CHIKV infection, it was further tested whether EG had any direct virucidal effect on CHIKV infectivity. After performing the virucidal assay, no significant difference in viral titers was observed in EG treated viral inoculum compared to the untreated infection control. (**Fig. S2H**).

**Figure 4:**
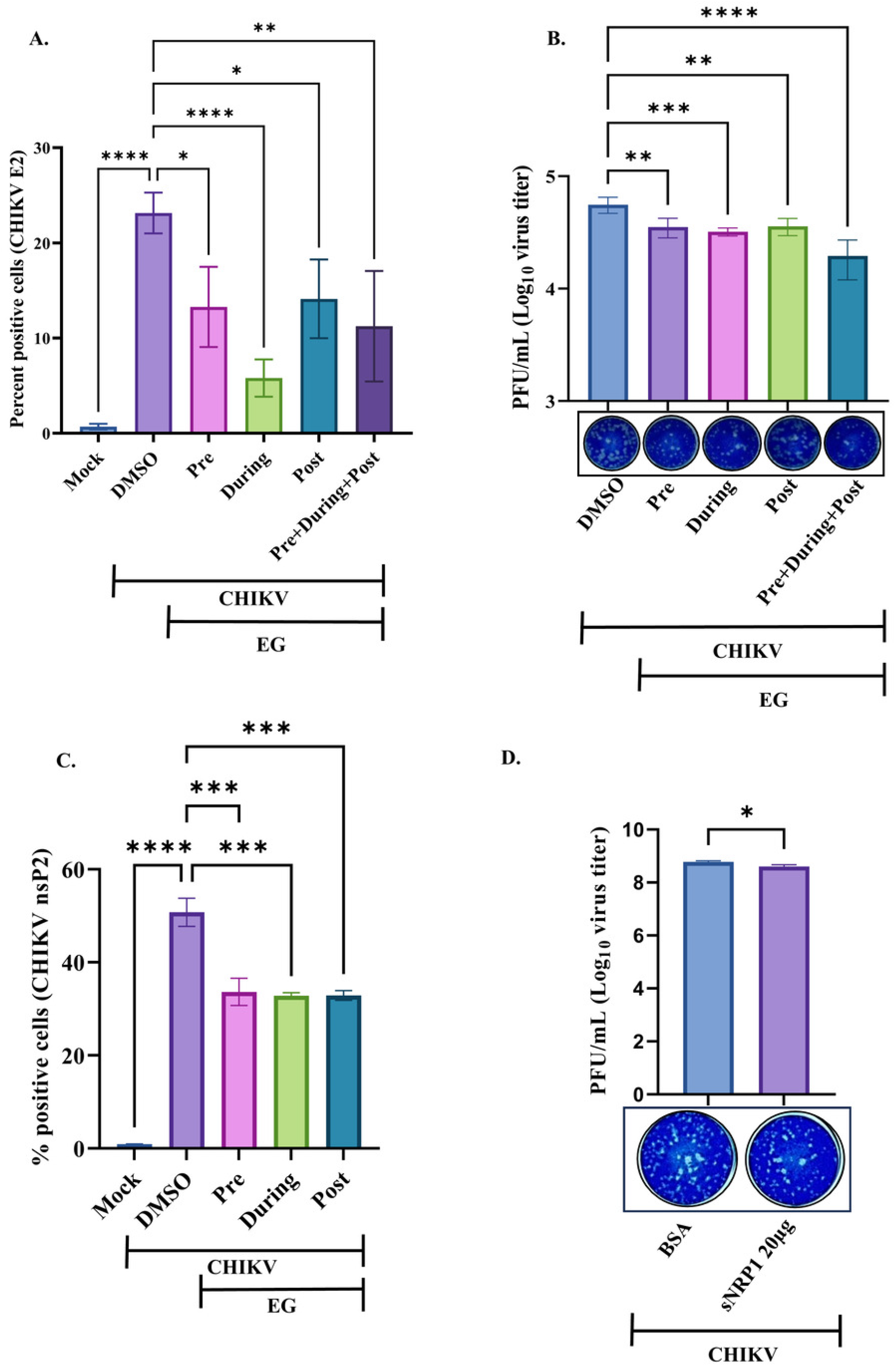
NRP1 is involved in multiple phases of CHIKV infection. **(A**) Bar graph showing the percentage of CHIKV E2-positive RAW 264.7 cells following EG treatment at different phases of CHIKV infection, and **(B)** Bar graph showing viral titers upon EG treatment across different phases of CHIKV infection. **(C)** Bar graph representing % CHIKV nsP2 positive cells following EG treatment at different phases of infection in C2C12 cells. **(D)** Bar graph demonstrating reduced viral titers in soluble NRP1 blocking assay in C2C12 cells. Data are presented as the mean ± SD from three independent experiments. Statistical significance was determined as indicated (ns, not significant; *, P < 0.05; **, P ≤ 0.01; ***, P ≤ 0.001; ***, P ≤ 0.0001).

Since, EG treatment reduced viral titers both in -pre and -during phase of CHIKV infection, soluble NRP1 blocking assay was performed to better understand the requirement for NRP1 during the entry or attachment phase. CHIKV was incubated with soluble NRP1 (sNRP1) prior to infecting C2C12 cells. BSA was used as negative control. Post infection, plaque assay was performed to determine the viral titer at 15hpi. It was observed that there was reduction (approximately 37% decrease) in viral titer with sNRP1 treatment (**Fig. 4D**). Overall, these findings suggest that NRP1 might be involved in multiple phases of CHIKV infection.

### 2.6. NRP1 interacts with CHIKV envelope protein E1 during viral infection

Given the involvement of NRP1 in multiple stages of CHIKV infection, including viral attachment and entry, as demonstrated by the pre, during, and post-infection assays and the soluble NRP1 blocking assay, its interaction with the CHIKV envelope proteins was investigated. To address the same, protein-protein molecular docking was performed between NRP1 complex (PDB ID: 4GZ9 mouse, and 2QQM human) and CHIKV Envelope protein (PDB ID: 3N41) **(Fig. 5A-B).** The analysis showed 9 and 15 probable interactions between the amino acid residues of human NRP1-CHIKV E1 and mouse NRP1-CHIKV E1 respectively (**Table S1-S2**). This suggests a possibility of interaction of CHIKV-E1 with the b1b2 domain of NRP1 that might be required for the efficient viral infection in host. *In silico* interactions were also found in NRP1-CHIKV E2 complexes, however; NRP1-CHIKV E1 complex appeared to be stronger and more biologically relevant (**Fig. S3A-B**, **Table S1-S2**).

**Figure 5:**
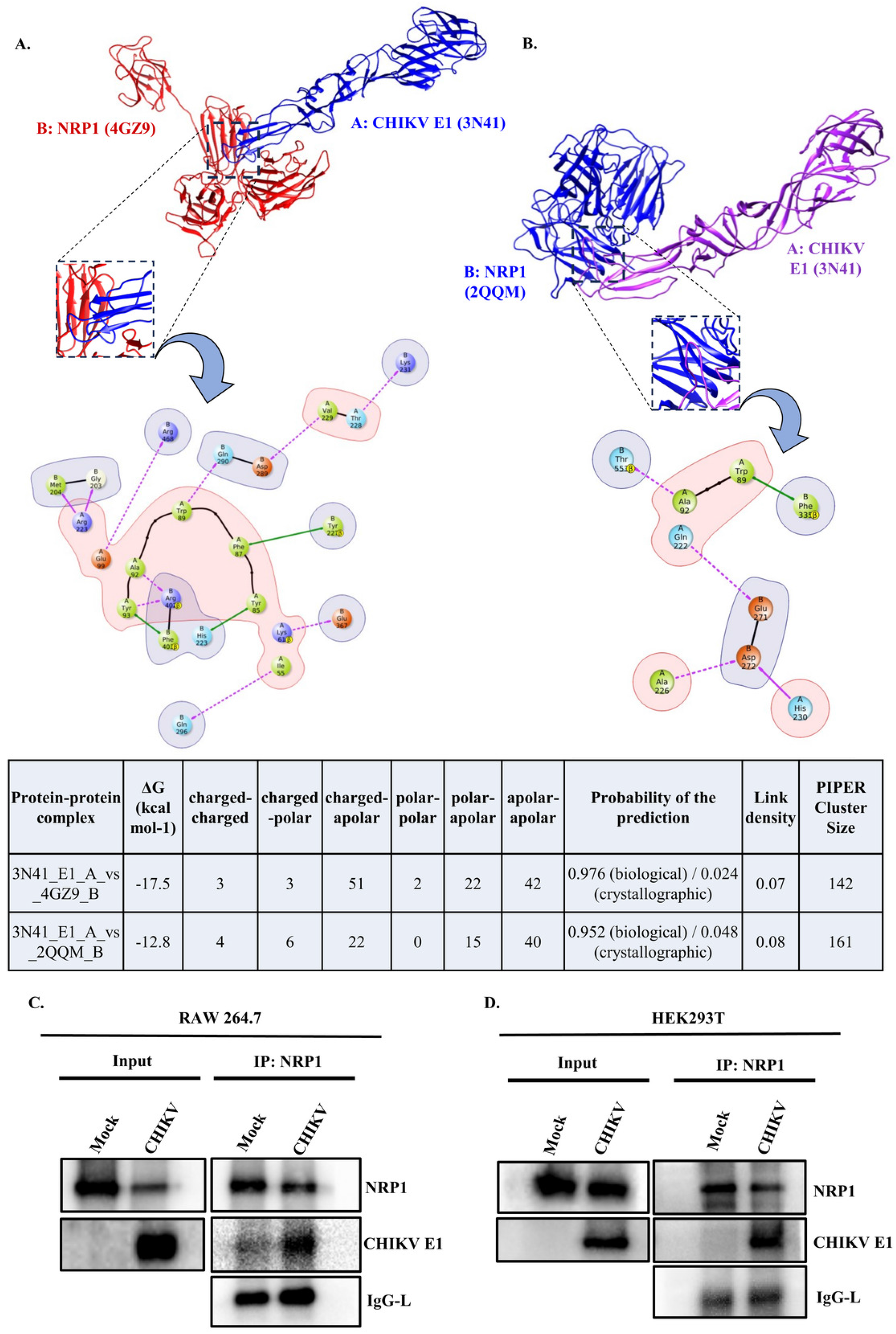
NRP1 interacts with the CHIKV envelope protein E1. Molecular docking was performed between NRP1 and CHIKV envelope proteins using Schrödinger’s Maestro. For *in vitro* interaction studies, the RAW 264.7 and HEK cells were infected with CHIKV and processed for immunoprecipitation followed by Western blot analysis. Protein-protein docking analysis depicting best docking pose and 2D interactions of **(A)** mouse NRP1 (PDB ID: 4GZ9) with CHIKV E1 (PDB ID1: 3N41), and **(B)** human NRP1 (PDB ID: 2QQM) with CHIKV E1 (PDB ID1: 3N41), and comparative table showing nature and strength of the interactions. Western blot analysis showing the expressions of E1 and NRP1 in the input (whole cell lysate: left), and co-immunoprecipitated lysate depicting the interaction of CHIKV E1 and NRP1 (right) in **(C)** RAW 264.7 cells, and in **(D)** HEK cells.

To validate the *in-silico* results, co-immunoprecipitation (co-IP) assay was performed. RAW 264.7 and HEK cells were infected with CHIKV at MOI 5 (for 8hpi), and 0.1 (for 15hpi) respectively. Infected cells were harvested and subjected to co-IP assay followed by Western blot analysis. It was found that, CHIKV-E1 immunoprecipitated with host NRP1 in both RAW 264.7 and HEK cells (**Fig. 5C-D**). Next, to determine whether the host NRP1 protein interacted with CHIKV E2, similar experiments were performed to pulldown NRP1 with CHIKV E2 (**Fig. S3C**). However, NRP1 was not detected in the pull-down fraction of CHIKV E2. Collectively, these results indicate that NRP1 interacts with CHIKV envelope protein E1.

### 2.7. NRP1 antagonism augments macrophage activation during LPS treatment and CHIKV infection in macrophages

Lipopolysaccharide (LPS) induced experimental immune activation model was used to determine the inherent activity of EG mediated NRP1 inhibition on macrophage activation and polarization. The RAW 264.7 cells were treated with EG (10µM) in presence of LPS (200ng/mL) for 10 hours; and the surface expression of CD86, MHC I/II, CD64 (M1 polarization marker), NRP1, and CD206 (M2 polarization marker) was evaluated by flow cytometry (**Fig. S4A**). It was observed that LPS treatment downregulated NRP1 (**Fig. S4B**). EG treatment increased the previously downregulated NRP1 expression during LPS treatment (**Fig. S4F**). On the other hand, EG treatment alone led to a marked upregulation of MHC I, and NRP1; and a modest increase in MHC II, CD86, and CD64. In addition, LPS+EG treatment significantly upregulated MHC I, NRP1, and CD64; while a modest upregulation of MHC II, CD86, and CD206 was seen as compared to only LPS treated cells (**Fig. S4C-G**).

The effects of EG mediated NRP1 inhibition on macrophage activation and polarization during CHIKV infection were further analyzed. Cells were treated with EG during CHIKV infection and the surface expression of MHC II, CD86, CD64, CD206 was quantified by flow cytometry (**Fig. S5A**). CHIKV infection significantly increased the expression of MHC II, CD86, CD64, and CD206 compared with mock. While EG treatment led to a marked increase in MHC II expression; a modest increase in CD86, and CD64 expression was observed as compared to only CHIKV infected cells (**Fig. S5D).** However, EG treatment did not significantly alter the expression of the M2 macrophage polarization marker CD206 compared with CHIKV-infected cells. **(Fig. S5E)**. These results suggest that NRP1 inhibition might augment activation of macrophages during LPS-treatment and CHIKV-infection.

### 2.8. NRP1 inhibition leads to enhanced JNK signaling and heightened inflammatory cytokine responses during CHIKV infection

To assess the effect of NRP1 inhibition upon MAPK signaling proteins during CHIKV infection, phosphorylation of SAPK/JNK and p38 MAPKs was analyzed by Western blot (**Fig. 6A**). EG mediated NRP1 inhibition markedly decreased CHIKV E2 (as previously shown in Fig. 1J) while increasing phosphorylation of SAPK/JNK in CHIKV-infected RAW 264.7 cells, with no significant change in p38 phosphorylation (**Fig. 6B-C)**. Notably, anti-NRP1 antibody treatment also decreased CHIKV E2 (as previously shown in Fig. 1J), and increased p-SAPK/JNK protein levels, without significantly changing of p-p38 levels, consistent with the EG-mediated effect (**Fig. 6D-F**).

**Figure 6:**
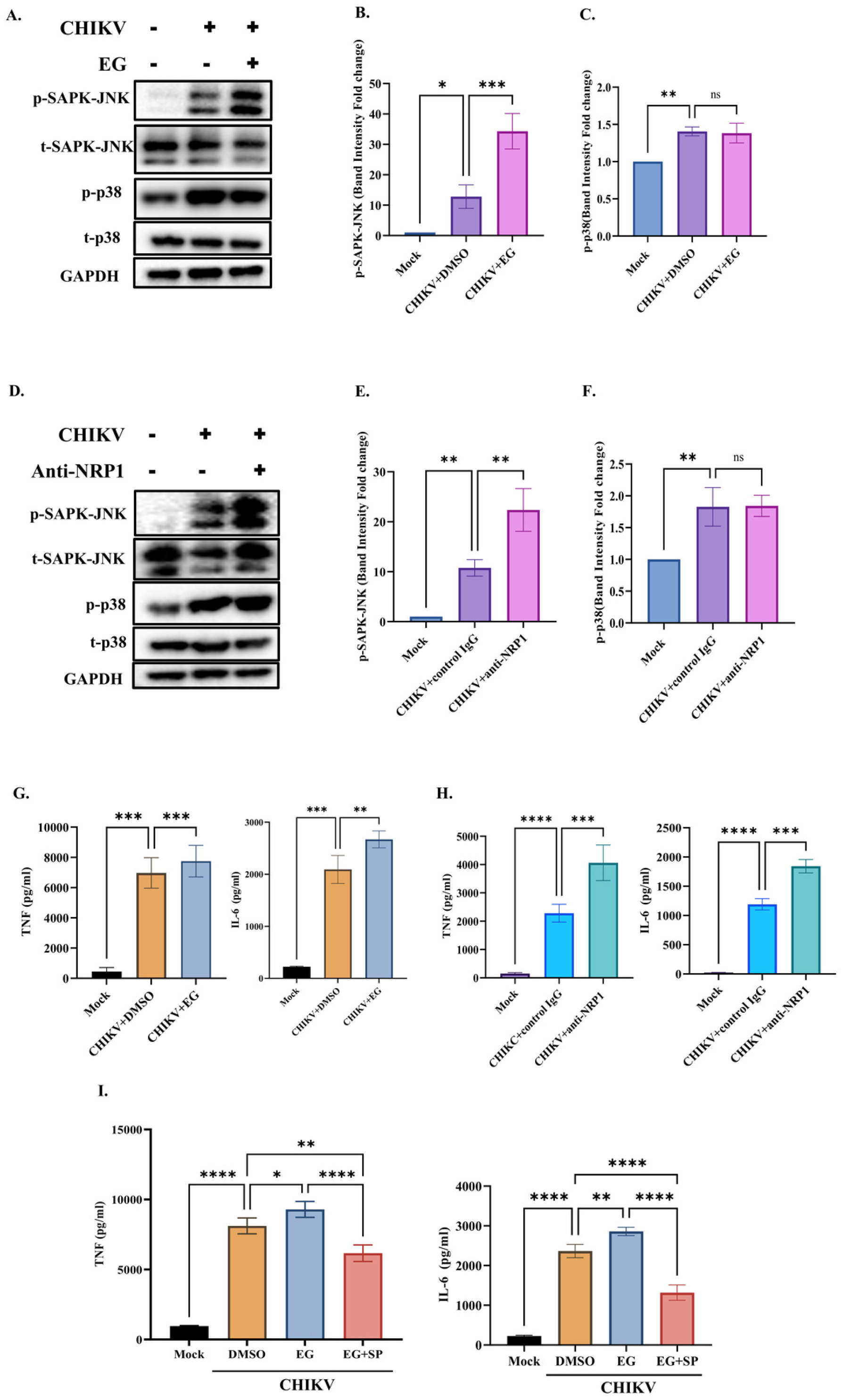
NRP1 inhibition enhances JNK signaling and inflammatory cytokine responses during CHIKV infection. RAW 264.7 cells were treated with DMSO or EG00229 trifluoroacetate (EG). For antibody-mediated blockade, cells were treated with anti-NRP1 or isotype control IgG antibodies. Cells were infected with CHIKV at an infection multiplicity (MOI) of 5 for 2 h and harvested at 8 hpi. **(A)** Immunoblot analysis showing differential expression phosphorylated and total SAPK/JNK, and p38 MAPK following EG-mediated NRP1 inhibition in CHIKV-infected RAW 264.7 cells. Densitometry analysis and fold change (with respect to mock) of **(B)** p-SAPK-JNK, **(C)** p-p38. (**D)** Immunoblot analysis showing the expression of phosphorylated and total SAPK/JNK, and p38 MAPK in NRP1 antibody blockade-mediated inhibition in CHIKV-infected RAW 264.7 cells. Densitometry analysis and fold change (with respect to the mock) of **(E)** p-SAPK-JNK, **(F)** p-p38 due to NRP1 antibody blockade. **(G)** ELISA-based quantification of TNF-α and IL-6 levels during EG-mediated NRP1 inhibition, and **(H)** anti-NRP1 antibody blockade. **(I)** ELISA-based quantification of TNF-α and IL-6 levels in CHIKV-infected RAW cells treated with EG+SP co-treatment. Data are presented as the mean ± SD from three independent experiments. Statistical significance was determined as indicated (ns, not significant; *, P < 0.05; **, P ≤ 0.01; ***, P ≤ 0.001; ***, P ≤ 0.0001).

To further elucidate the downstream effects of NRP1 inhibition on inflammatory cytokine response in CHIKV-infected macrophages, secreted TNF and IL-6 levels were quantified by ELISA. Both EG treatment and anti-NRP1 blockade significantly increased TNF and IL-6 secretion compared to CHIKV infection alone **(Fig. 6G-H)**.

To further investigate whether the enhanced inflammatory cytokine response elicited by NRP1 inhibition was JNK mediated, dual inhibition of NRP1 and JNK was performed using their selective inhibitors (EG and SP, respectively). A combinatorial treatment of 10µM of EG along with 10µM of SP had no cytotoxic effect on cell viability (**Fig. S5F**). Co-treatment of SP and EG mitigated the CHIKV-induced reduction in NRP1 expression (**Fig. S5G**). Interestingly, SP mediated JNK inhibition significantly suppressed the EG-induced elevation of TNF and IL-6 in infected RAW 264.7 cells (**Fig. 6I**). In addition, simultaneous inhibition of NRP1 and JNK had an additive anti-viral effect on CHIKV infection, as evidenced by further reduction of % CHIKV E2 macrophages **(Fig. S5H).** Together, these results show that NRP1 inhibition may promote a JNK-driven inflammatory cytokine response in CHIKV-infected macrophages.

### 2.9. EG reduces viral load and elicits protective cytokine response in CHIKV infected mice

The antiviral efficacy of EG was evaluated in 10-12-days old C57BL/6 mice that were subcutaneously infected with 10⁶ PFU of CHIKV. Mice received oral dosage of EG (5 mg/kg) at every 24 h intervals, starting from 2 days prior to infection until 1 day post infection (dpi) (**Fig. 7A**). At 5 dpi, mice were sacrificed followed by downstream experiments. EG treatment significantly reduced viral burden in both brain and muscle (more than 90%) (**Fig. 7B-C**) compared to only infected. The infected mice showed more arthritis in their limbs and greater mobility impairment than the treated mice (**Fig. 7D**). Consistently, reduction in protein levels of CHIKV E2 (∼81.7% reduction) and nsP2 (∼52.9% reduction) were also observed in the brain of EG-treated mice (**Fig. 7E-F**). Similarly, Western blot analysis revealed markedly lower levels of CHIKV E2 (∼63.2% reduction) and nsP2 (∼61.9% reduction) proteins in muscle tissue following EG treatment (**Fig. 7G-H)**. As seen in the *in vitro* observations, NRP1 protein expression was downregulated in both brain and muscle tissues in infected mice. EG treatment did not alter NRP1 expression in the brain but reduced NRP1 expression in the muscle tissues as compared to CHIKV-infected mice **(Fig. 7F, 7H**). While the infected group exhibited weight loss from 3 dpi, the average body weight of the EG-treated group of mice was comparatively stable (**Fig. 7I**). Interestingly, serum cytokine analysis demonstrated significantly elevated levels of IFN-γ, and IL-6, accompanied by reduced IL-17A levels, in EG-treated mice compared to the only infected mice (**Fig. 7J**). Collectively, these findings suggest that EG00229 trifluoroacetate exhibits anti-CHIKV efficacy in a mouse model by reducing viral burden and modulating host immune responses.

**Figure 7:**
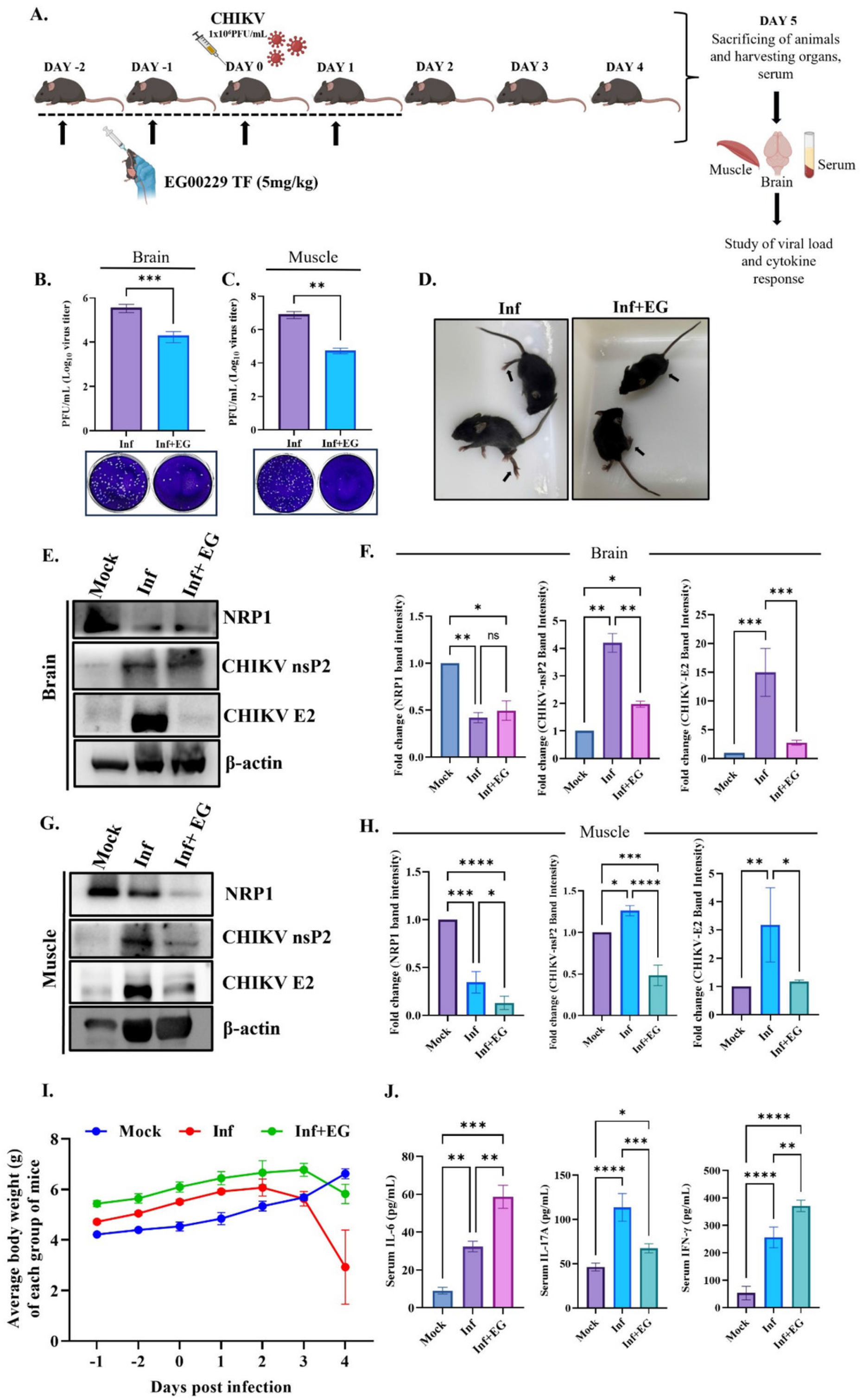
EG Treatment induces anti-viral immune response in CHIKV infected mice model. C57BL/6 mice were subjected to EG00229 Trifluoroacetate (EG) at dosage 5mg/kg for 2 days prior to subcutaneous infection with 10^6^ PFU of CHIKV. They were further treated with EG up to 1 dpi. Mice were sacrificed at 5 dpi, and serum and different tissues were collected for further downstream experiments. **(A)** Infection and EG treatment regimen in *in vivo* mice model of infection. Bar graph depicting viral load in **(B)** brain, and **(C)** muscle in CHIKV infected mice with EG treatment. **(D)** Image of CHIKV-infected and drug-treated mice. **(E)** Representative Western blot images, and **(F)** bar graphs showing protein levels of NRP1, CHIKV E2 and CHIKV nsP2 in brain tissue. **(G)** Representative Western blot images, and **(H)** bar graphs showing protein levels of NRP1, CHIKV E2 and CHIKV nsP2 in muscle tissue. Actin was used as loading control. Fold change shown with respect to mock control. **(I)** Average body weight of each group of mice. **(J)** Bar diagrams indicating the concentrations (pg/ml) of different cytokines (IL-6, IL-17A, IFN-γ) in serum of mock, infected, and EG-treated mice. Data are presented as the mean ± SD (body weight as Mean+SEM) from three independent experiments. Statistical significance was determined as indicated (ns, not significant; *, P < 0.05; **, P ≤ 0.01; ***, P ≤ 0.001; ***, P ≤ 0.0001).

### 2.10. Pharmacological inhibition of NRP1 decreases virus infection in hPBMC-derived monocyte-macrophage populations, and augments proinflammatory cytokine response, *in vitro*

To study the effect of NRP1-mediated regulation of CHIKV infection in a higher-order mammalian system, hPBMC derived adherent macrophage population (97% CD14^+^CD11b^+^ cells) (**Fig. 8A**) was subjected to infection in the presence and absence of EG (10μM) as per protocol mentioned previously (5, 33, 34). 10µM of EG was found to be non-cytotoxic dosage as per MTT assay (**Fig. S1H**). The hPBMC-derived adherent populations collected from 3 healthy donors showed a decrease in the percentage of CHIKV E2-positive cells (∼35% reduction) and in the mean fluorescence intensity (MFI) of CHIKV E2 (∼45.3% reduction) with EG treatment (**Fig. 8B-D**). In addition, there was also a reduction in CHIKV titer after EG treatment as observed by the plaque assay (∼2.092-fold reduction) (**Fig. 8E**). To assess the effect on cytokine responses, the secreted TNF level was determined by ELISA. Significantly higher TNF secretion was observed in EG-treated cells as compared to only CHIKV (**Fig. 8F**). Collectively, these data indicate that NRP1 antagonism reduces CHIKV infection and upregulates pro-inflammatory response in the hPBMC-derived monocyte/macrophages.

**Figure 8:**
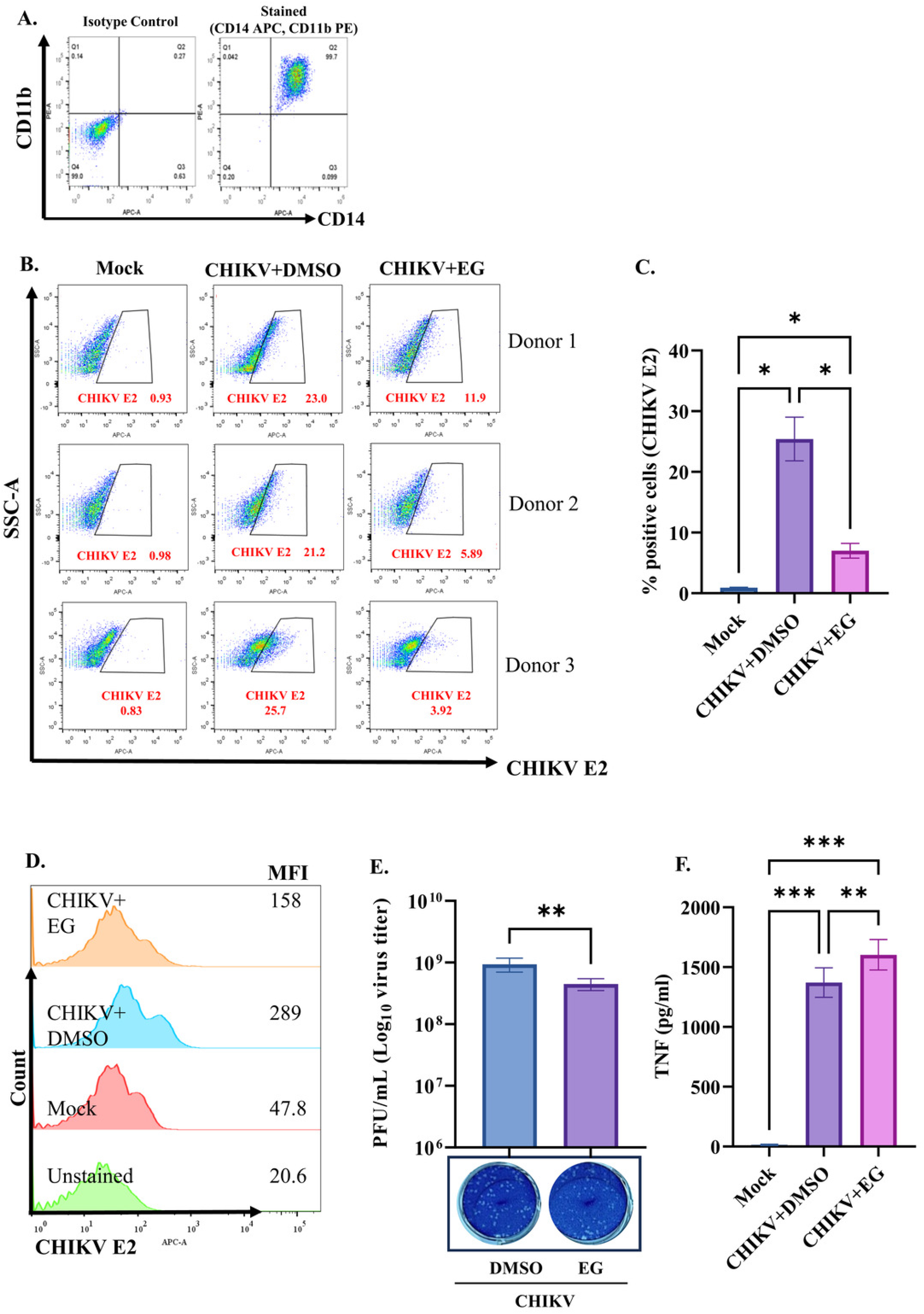
Pharmacological inhibition of NRP1 decreases virus infection in hPBMC-derived monocyte-macrophage populations and augments the proinflammatory cytokine response. The hPBMC-derived adherent monocyte-macrophage cells were pretreated with DMSO or EG for 2 h prior to CHIKV infection. The CHIKV infection was given at 5 MOI for 2 hours and cells were harvested at 8 hpi. **(A**) Flow cytometry-based immunophenotyping of adherent populations of hPBMC-derived myeloid cells, showing that 99% of the cells were CD14+ CD11b+. **(B)** Representative flow cytometry dot plots, and **(C)** bar graph showing the %CHIKV E2-positive cells with EG treatment in CHIKV infected hPBMCs. **(D)** Mean fluorescence intensity (MFI) analysis of CHIKV E2 expression. **(E)** Bar graph showing CHIKV titers. **(F)** ELISA-based quantification of human TNF levels in culture supernatants. Data are presented as the mean ± SD from three independent experiments. Statistical significance was determined as indicated (ns, not significant; *P < 0.05; **P ≤ 0.01; ***P ≤ 0.001; ****P ≤ 0.0001).

## 3. DISCUSSION

CHIKV pathogenesis involves a fine balance between viral infection and host immune responses. Thus, host factors that control immune homeostasis, are critical determinants of viral control and immunopathology (7). This study identifies a previously unrecognized role for Neuropilin-1 (NRP1) in CHIKV pathogenesis as a host factor that promotes viral infection and regulates host immune responses.

In this study, we have identified NRP1 as an important host factor in CHIKV infection. Inhibition of NRP1 (by EG treatment, antibody blocking, and siRNA knockdown) leads to a significant reduction in viral burden, whereas NRP1 overexpression enhances CHIKV infection. Moreover, NRP1 is involved in multiple phases of CHIKV infection and interacts with the viral envelope protein E1. In addition, NRP1 regulates the inflammatory JNK pathway in macrophages, thereby shaping cytokine responses during infection. Furthermore, the anti-CHIKV and immunomodulatory effects of the NRP1 inhibitor (EG) were validated in human primary cells, and in the murine model of CHIKV infection. These findings suggest that NRP1 contributes to efficient viral infection and immune regulation during CHIKV pathogenesis.

Several studies have reported that NRP1 facilitates the infection of various viruses like SARS-CoV-2, murine cytomegalovirus, Kaposi sarcoma virus (KSHV), Epstein-Barr Virus (EBV), Human T-cell Leukemia Virus-1 (HTLV-1), Mammalian orthoreovirus, Enterovirus A71 (EVA71), hepatitis B virus (HBV), and Pseudorabies virus (PRV) (25–30, 35). The current study explores the contribution of NRP1 in alphavirus infection by using Chikungunya virus as a model system. NRP1 expression was found to be downregulated in CHIKV infected macrophages. This observation aligned with previous reports in which NRP1 downregulation was associated with viral infection and heightened inflammatory response (15, 20, 29). EG00229 trifluoroacetate (EG) is a potent functional inhibitor of NRP1; that binds to the extracellular region of NRP1, preventing it from binding to its co-receptors and ligands (36). Thus, as expected, EG treatment did not significantly alter the surface NRP1 expression in macrophages, irrespective of their infection status. EG has been previously shown to reduce SARS-CoV-2, and HBV infections (27, 36). Likewise, our current study demonstrates that treatment with EG, reduced CHIKV load and infectious titers in macrophages. Furthermore, EG treatment exhibited anti-CHIKV activity in several cell types, including muscle, microglial, and epithelial cells. In addition, siRNA-mediated knockdown of NRP1 reduced viral protein expression and infectious titers, whereas NRP1 overexpression increased viral load. Our observations aligned with previous studies in other viruses such as HTLV-1, Orthoreovirus, SARS-CoV-2, and EBV; where NRP1 expression was positively correlated with virus infection (25–27, 30, 35).

Previous reports have extensively highlighted the role of NRP1 in mediating attachment and entry of several viruses (25–29, 35). In the current study, antibody blocking of membrane NRP1 using monoclonal anti-NRP1 antibodies, led to reduced viral titers and viral protein levels in macrophages, similar to previous studies in SARS-CoV-2 (25). Furthermore, pre-incubation of the viral inoculum with soluble NRP1 resulted in reduced viral titers. This underscores a probable role of host NRP1 in the early entry phase of CHIKV infection, similar to a previous study done with KSHV (28). Interestingly, EG treatment reduced viral titers across all stages of CHIKV infection including pre, during infection, and post-viral attachment phases. In addition, it was also observed that EG did not have a direct effect on CHIKV infectivity, and most likely exerts anti-viral effects by modulating the host NRP1. Altogether these suggest the possible pro-viral role of NRP1 across multiple phases of the CHIKV life cycle, extending beyond viral entry. However, further detailed studies are required to elucidate the detailed underlying mechanisms.

Direct interactions between NRP1 and viral proteins have been experimentally demonstrated to facilitate infection across multiple viruses. NRP1 was demonstrated to interact with SARS-CoV-2 spike protein, enterovirus VP3 capsid protein, pseudorabies virus glycoproteins, EBV glycoprotein B (gB), HTLV-1 Env, KSHV glycoprotein B (gB), and HBV preS1 (26–30, 36, 37). In the current study, *in silico* molecular docking revealed specific amino acid residues interacting between host NRP1 and CHIKV E1. While NRP1 and E1 significantly showed maximum biologically relevant interaction, potential interacting residues were also observed between NRP1 and E2. The higher link density and biological relevance of the E1-NRP1 complexes may be attributed to a more favourable balance of apolar and polar interactions compared with the corresponding E2-NRP1 complexes. To validate the interaction *in vitro;* CHIKV infection followed by co-immunoprecipitation was performed between CHIKV envelope proteins and NRP1. Co-IP studies demonstrated that, NRP1 interacts with CHIKV-E1 protein in CHIKV-infected RAW 264.7 and HEK cells. NRP1 effectively immunoprecipitated the CHIKV-E1 protein. However, NRP1 did not immunoprecipitate with CHIKV E2 *in vitro*; demonstrating the specificity of the interaction between NRP1 and E1. Thus, the pro-viral nature of NRP1 might be attributed, at least in part to its favourable interaction with CHIKV envelope protein E1, which requires further in-depth studies.

Beyond facilitating viral entry, NRP1 has been linked with immunoregulation in diverse pathological conditions, including sepsis, autoimmunity, cancer, and chronic inflammatory diseases, and has been implicated in promoting anti-inflammatory M2 macrophage polarization (14). In the current study, RAW 264.7 macrophages exhibit increased expression of MHCs, CD86, and CD64 (M1 marker) during LPS treatment and CHIKV infection as reported earlier (5, 38). Interestingly, NRP1 inhibition further elevated CD86, MHC II, and CD64 expression with no significant change in CD206 (an M2 marker) during CHIKV infection. This suggests that NRP1 antagonism might lead to a more immune-activated macrophage phenotype associated with antiviral immunity.

The macrophage response to CHIKV infection is generally associated with elevated MAPK signalling, particularly increased phosphorylation of JNK and p38 (5, 39, 40) . Moreover, enhanced JNK activation has been linked to pro-inflammatory cytokine responses in CHIKV and other diseases (39, 40). In the current study; NRP1 inhibition in RAW 264.7 macrophages increased JNK phosphorylation without significantly affecting p38 activation. Increased JNK activation upon NRP1 blockade suggests that NRP1 might suppress JNK-driven inflammatory responses during CHIKV infection. As reported previously, CHIKV infection leads to the enhanced production of inflammatory cytokines, including TNF and IL-6, (5, 41). Interestingly, NRP1 inhibition in infected macrophages led to further elevated levels of pro-inflammatory cytokines (TNF, and IL-6). Our observation aligns with several previous reports, in which the absence of NRP1 led to a heightened response in inflammatory diseases (15, 16). To further assess whether NRP1 antagonism leads to an elevated cytokine response mediated via JNK signalling, a selective JNK inhibitor, SP; was used. Notably, JNK inhibition attenuated the EG-mediated elevation of pro-inflammatory cytokines, suggesting that NRP1-dependent immunomodulation may be at least be partially JNK-dependent. Interestingly, co-treatment with SP+EG also counteracted the previously downregulated NRP1 expression due to CHIKV-infection, linking the functional NRP1 expression to immune activation and inflammatory cytokine responses. Moreover, concurrent inhibition of NRP1 and JNK resulted in a greater reduction in CHIKV infection. Since JNK signalling is a key regulator of stress responses, cytokine production, and inflammatory programming in CHIKV-infected macrophages (32, 40), the enhanced antiviral effect observed with dual inhibition suggests that NRP1 may either directly modulate JNK activation or converge on shared downstream effectors that may together create a proviral cellular environment. In addition, NRP1 appears to regulate macrophage immune homeostasis during CHIKV infection. Simultaneous inhibition of NRP1 and JNK likely perturbs this immunoregulatory balance even more, thereby limiting viral infection. Our observations suggest that this further enhancement of M1 phenotype and pro-inflammatory responses in macrophages during acute CHIKV infection might help in the better resolution of viremia. The precise mechanism however requires further investigation.

The *in vitro* anti-viral and immunomodulatory roles of the NRP1 inhibitor were further explored in a C57BL/6 mouse model. There was a significant reduction in CHIKV viral titer, and viral protein levels in EG-treated CHIKV-infected C57BL/6 mice. These outcomes are consistent with previous reports demonstrating anti-viral efficacy of EG against HBV infection in mice (27). Interestingly, this decline in viral load was accompanied by an elevated IL-6, IFN-γ and reduced IL-17A secretion. While IFN-γ is a hallmark for Th1 response, IL-6 is a pleotropic cytokine that also supports maintenance of effector T-cell responses during antiviral immunity (42, 43). Moreover, IFN-γ programs Tregs into Th1-like phenotype, and controls virus-specific T-cell immunity during viral infection (44). Therefore, the elevated IFN-γ and IL-6 observed following EG treatment likely reflects a more prominent antiviral immune environment. In addition, CHIKV is often associated with the expansion of regulatory RORγt⁺ Th17 cells that secrete IL-17A (45). IL-17A is an inflammatory cytokine, that also promotes Th17 Treg-mediated immunosuppression. Moreover, IL-17A-deficient mice exhibit increased IFN production, reduced tissue inflammation, and attenuated joint swelling following CHIKV infection (46). Notably, NRP1 is reported to be a positive regulator of IL-17A (47). Collectively, it can be suggested that NRP1 can influence IL-17A-mediated responses during CHIKV infection. Interestingly, our observations indicate that EG-mediated NRP1 inhibition might subdue IL-17A-dependent inflammation while enhancing Th1 responses. This aligns with our *in vitro* studies where EG led to an anti-viral response with upregulated M1 phenotype in infected macrophages. These findings suggest that NRP1 inhibition might differentially reprogram the immune response towards a more effective anti-viral phenotype while also contextually limiting IL-17A driven immunopathology. This balanced immune activation might appear to facilitate viral clearance, supporting a protective role for EG-induced immunomodulation in reducing CHIKV burden.

Human blood derived PBMCs have been previously used as models to study the implications of human innate immune responses and to evaluate the efficacy of antiviral and host-directed therapeutic intervention (5, 48–50). Thus, the anti-viral potential of the NRP1 inhibitor was further explored in human primary PBMCs. Notably, NRP1 inhibition using EG reduced infection and viral titers in the monocytes/macrophages isolated from PBMCs of healthy human blood donors. The hPBMCs secrete TNF in response to viral infection as mentioned in previous studies (5, 50). EG treatment led to an increased TNF response consistent with our *in vitro* observations in macrophage cell lines. Thus, NRP1-dependent modulation of CHIKV infection and inflammatory responses extended to primary human immune cells, underscoring its physiological relevance in higher-order mammalian systems.

Our findings identify NRP1 as a previously unrecognized proviral and immunoregulatory host factor that interacts with CHIKV structural protein, influences multiple stages of CHIKV infection, and shapes antiviral immune responses *in vitro* and *in vivo*. However, the precise mechanism governing the anti-CHIKV activity of NRP1 inhibition remains to be further explored using knockout cells and conditional NRP1 deficient mice. Further studies could also be extended to additional infection models, such as hamster, zebra fish, and ultimately validated in clinical samples of CHIKV-infected patients. In addition, while the NRP1 inhibitor enhances antiviral immune activation during CHIKV infection, its translational potential could be strategically augmented by combining it with anti-inflammatory immune modulators, which requires further investigation.

Collectively, the current study established for the first time NRP1 as a multifunctional host factor during CHIKV pathogenesis, coordinating infection dynamics and viral immune response. Thus, targeting NRP1 may represent an effective host-directed approach to control CHIKV infection while positively modulating immune responses.

## 4. MATERIAL AND METHODS

### 4.1. Cells, and Virus

The RAW 264.7, THP-1, and BV2 cells were cultured and maintained in RPMI-1640 medium (Gibco, USA) supplemented with penicillin-streptomycin antibiotic solution (HiMedia, India) and 10% heat-inactivated fetal bovine serum (FBS; Gibco, USA) at 37⁰C in a humidified incubator with 5% CO_2_. Vero and HEK293T cells were maintained in DMEM media (Gibco, USA) supplemented with penicillin-streptomycin antibiotic solution and 10% heat-inactivated FBS at 37⁰C in a humidified incubator with 5% CO2. C2C12 cells were grown in DMEM under the same conditions with 15% FBS. The CHIKV prototype strain (PS, accession no. AF369024.2), and anti-CHIKV E2 monoclonal antibody were kindly provided by Dr. M.M. Parida, (DRDE, Gwalior, India). The Indian outbreak strain of CHIKV (IS; accession no. PP349434) was also used in this study. CHIKV-IS was used for *in vitro* infection studies, and CHIKV-PS strain was used for the *in vivo* work. The CHIKV nsP2 antibody used in this study was developed by us (51). RAW 264.7, THP-1, and C2C12 were obtained from American Type Culture Collection (ATCC), USA. Vero and HEK293T were purchased from National Centre for Cell Science (NCCS), Pune, India. C2C12 cells were a kind gift from Dr. Amaresh C. Panda (Institute of Life Sciences, Bhubaneswar), and BV2 cells were kindly provided by Dr. Anirban Basu (National Brain Research Center, India). All the experimental protocols involving CHIKV were conducted in accordance with the Institutional Biosafety Committee (IBSC) guidelines.

### 4.2. Antibodies and Reagents

EG00229 Trifluoroacetate (Cat. 6986), a potent NRP1 inhibitor, was purchased from Tocris Biosciences, UK. The antibodies against NRP1 (Fluorochrome PE-Cy7, Cat. 25-3041-82), CD86 (Fluorochrome: APC; Cat. 17-0862-82), and MHC-II (Fluorochrome: PE; Cat. 12-5321-82) were purchased from eBiosciences, USA. The antibodies used for western blot: total p38 (Cat. 9212), phosphorylated p38 (Cat. 9211), total SAPK-JNK (Cat. 9252), and phosphorylated SAPK-JNK (Cat. 4668), were purchased from Cell Signaling Technology (Denver, USA). The rabbit NRP1 monoclonal antibody (Cat. ab81321, clone: EPR3113, abcam, USA), and rabbit CHIKV-E1 monoclonal antibody (GTX637973, GeneTex, USA) were used for Western blot and co-immunoprecipitation studies. The VeriBlot IP detection reagent (HRP) used in co-immunoprecipitation study (Cat. ab131366) was purchased from Abcam, USA. Fluorochrome (AF488/AF647) conjugated anti-mouse and rabbit secondary antibodies (used for flow cytometry) and HRP-conjugated anti-mouse and rabbit secondary antibodies (used for Western Blot) were purchased from Invitrogen, USA. Dynabeads protein A (10006D) and protein G (10007D) immunoprecipitation kits were bought from Thermo Fisher Scientific, USA. The GAPDH antibody (Cat. 10-10011) was bought from Abgenex India Pvt. Ltd, Bhubaneswar, India. Lipopolysaccharide (L4391-1MG), saponin, bradford reagent, NP-40, sodium deoxycholate, sodium dodecyl sulfate (SDS), protease inhibitor cocktail, and bovine serum albumin fraction V (BSA) were purchased from Sigma-Aldrich. The BCA reagent was bought from Invitrogen, USA. The Nitrocellulose membrane, PVDF membrane, and Immobilon Western Chemiluminescent HRP substrate were purchased from Millipore, USA. The ELISA kits-(mouse TNF (Cat. 558534), IL-6 (Cat. 555240), and human TNF (Cat. 55212), were bought from Waters Biosciences, USA. The mouse Th1/Th2/Th17 CBA Kit (Cat. 560485) was bought from Waters Biosciences, USA. The JNK inhibitor (SP600125) was from Merck. Recombinant mouse soluble NRP1(Cat. 5994-N1) was purchased from R&D systems, USA.

### 4.3. Cell Viability Assay

The working concentration of EG00229 trifluoroacetate (EG) in different cell lines (RAW 264.7, THP-1, Vero, BV2, HEK293T, and C2C12) and human peripheral blood mononuclear cells was determined using the MTT assay, according to the manufacturer’s protocol (Cat. CCK-003-2500, HiMedia, India). The non-cytotoxic dose of combined EG and SP treatment on RAW 264.7 cells was also determined using the MTT assay as described previously (5). The absorbance was measured at 560 nm using a microplate reader (BioTek, Epoch 2), and the cell viability was calculated in percentage.

### 4.4. CHIKV infection

As reported previously, RAW 264.7 cells were infected with CHIKV (5). Briefly, the cells were seeded into culture plates for 16-19 h prior to viral infection. The cells were then washed twice with 1X PBS before adding the virus inoculum at an MOI of 5 and incubated at 37 °C for 2 h. After 2 h, the cells were washed to remove unbound virus particles and incubated for 8-10 h (hours post-infection-hpi) in complete RPMI-1640 medium. BV2 cells were similarly infected like that of RAW 264.7 cells. HEK293T and C2C12 cells were infected at an MOI of 0.1 and 0.05, respectively, for 1.5 h, followed by incubation in complete media up to 15 hpi, as previously described (34, 52). Vero cells were infected with CHIKV at an MOI (multiplicity of infection) of 0.1 and harvested at 12 hpi. THP-1 cells were differentiated with 50 ng/mL phorbol myristate ester (PMA) for 12 h, followed by incubation with PMA-free complete RPMI-1640 medium for another 12 h. Differentiated THP-1 (dTHP-1) cells were then infected at an MOI of 5 using the same procedure as that of RAW 264.7 cells and harvested at 10 hpi. MOIs and time points that yielded maximal infection without inducing excessive cytopathic effects were selected based on previously reported studies for each cell type (34, 53).

### 4.5. EG00229 Trifluoroacetate (EG) Treatment

EG00229 trifluoroacetate (EG) is a selective antagonist of Neuropilin-1 that targets the b1 domain of its extracellular region, inhibiting ligand binding and consequently attenuating NRP1-dependent downstream signaling (54–56). All cells (RAW 264.7, dTHP-1, Vero, BV2, C2C12, hPBMCs) were pre-treated with 10µM of EG for 2 h prior to CHIKV infection, a concentration determined to be non-cytotoxic by MTT cell viability assays. As mentioned previously, EG was also added during and after infection, in accordance with the experimental requirements. DMSO was used as the solvent control. For experiments with the JNK inhibitor, EG was added along with SP600125 (SP) before infection, during infection, and post-infection in RAW 264.7 cells.

### 4.6. Plaque Assay

The plaque assay was performed using Vero cells to assess the viral titer per the earlier protocol (5). Briefly, cell-free culture supernatants from CHIKV-infected RAW 264.7, HEK, Vero, C2C12, dTHP-1, and hPBMCs were used to infect the Vero cells. After infection, 2% methyl cellulose (Cat. M0387; Sigma-Aldrich, USA) overlay DMEM media (HiMedia, India) was added to the infected cells and incubated for 3-4 days until visible plaques were formed. Next, the cells were fixed with 8% formaldehyde (Cat. M0387; HiMedia, India) and stained with crystal violet (Cat. C6158, Sigma) to manually determine the plaque-forming units (PFU) under white light using a transilluminator (Vilber Lourmat, France).

### 4.7. EG treatment at different phases of viral infection

To investigate the effect of EG at specific stages of viral infection; RAW 264.7 or C2C12 cells were treated with EG (10µM) 2 h before infection, during infection (viral adsorption), or post infection with CHIKV as per the protocol mentioned earlier (5, 34, 57). The combined condition included the addition of EG at all phases (pre, during, and post). Post-infection, cell free supernatants were collected and subjected to plaque assay to estimate viral titers.

### 4.8. Virucidal Assay

To investigate if EG directly affects CHIKV infectivity, CHIKV viral inoculum (0.1 MOI) was incubated with 10µM of EG for 30 min at 37°C as mentioned previously (57). Then, plaque assay was performed with the untreated and EG-treated viral inoculum to determine the residual infectivity.

### 4.9. Soluble NRP1 blocking Assay

For the soluble NRP1 (sNRP1) protein blocking assay was performed as reported earlier (28). CHIKV was preincubated with sNRP1 protein at indicated concentration or bovine serum albumin (BSA; as a negative control) at 37°C for 1 h, and then the sNRP1-bound CHIKV was used to infect C2C12 cells. At 15 hpi, cell free supernatants were collected and viral titers were estimated by plaque assay.

### 4.10. Flow Cytometry

The expression of surface and intracellular markers was evaluated using flow cytometry as described previously (5). An FcR-blocking reagent was used prior to primary antibody incubation to prevent nonspecific binding of antibodies to the Fc receptors on macrophages (cat. 130-092-575; Miltenyi Biotec, Germany), according to the manufacturer’s protocol. For intracellular staining, cells were fixed using 4% PFA (HiMedia) for 10 min, then washed with 1X PBS. The cells were then permeabilized using permeabilization buffer (1X PBS, 1% BSA, 0.1% saponin, and 0.01% NaN3) for 15 min at room temperature (RT), followed by blocking with a blocking buffer (1% BSA in permeabilization buffer) for 30 min at RT. The cells were then treated with primary antibodies (CHIKV E2 and CHIKV nsP2) for 30 min and incubated with the appropriate fluorophore-conjugated secondary antibody (AF647-conjugated chicken anti-mouse IgG (H + L) secondary antibody for 30 min). Both primary and secondary antibodies were diluted in permeabilization buffer. Both surface and intracellular stained samples were acquired using a BD LSRFortessa flow cytometer and analyzed using the FlowJo software (Waters Biosciences, USA). The cells were gated to exclude debris (FSC-A vs. SSC-A), followed by singlet discrimination (FSC-A vs. FSC-H) of live cells. Respective antigen marker-positive cells were quantified by SSC-A vs. the appropriate fluorescence marker within the live singlet population based on fluorescence intensity relative to the unstained and mock-infected controls.

### 4.11. Enzyme-linked immunosorbent assay (ELISA)

Cell-free culture supernatants from different experimental conditions were subjected to cytokine quantification using the Waters Biosciences OptEIA™ Sandwich ELISA kit (Waters Biosciences, USA) as described previously according to the manufacturer’s instructions (32). Cytokine quantification of mouse TNF, IL-6, and human TNF was performed with respect to standard curves prepared using recombinant cytokines at different concentrations (pg/mL).

### 4.12. Cytokine Bead Array

To measure cytokine levels in mice serum, samples were subjected to Cytometric Bead Array (CBA) assay according to the manufacturer’s instructions, as described previously (58). The standards were serially diluted in labelled tubes to create a standard curve, with a negative control containing only the assay diluent. The capture beads were mixed, and aliquots were added to the assay tubes along with the cytokine standards and samples. After the detection reagent was added, the tubes were incubated for 2 h at room temperature in the dark. Following incubation, wash buffer was added, and the tubes were centrifuged to discard the supernatant. Finally, the beads were resuspended in the wash buffer and acquired immediately via the BD LSRFortessa™ Cell Analyzer and analyzed using the FlowJo™ software, per the manufacturer’s guidelines.

### 4.13. Human peripheral blood mononuclear cell isolation and CHIKV infection

Studies involving human participants were reviewed and approved by the Institutional Ethics Committee of NISER, Bhubaneswar (NISER/IEC/2022-04). Human blood was drawn from healthy donors and myeloid adherent cells were generated from human peripheral blood mononuclear cells (hPBMC) as described previously (5). Briefly, circulating monocytes were enriched for 2 h of adherence after Hi-Sep LSM-based density gradient centrifugation according to the manufacturer’s instructions (HiMedia). The adherent cells were cultured in complete RPMI-1640 medium for 3-5 days. The adherent cells obtained after 96 h were of monocyte-macrophage lineage (more than 97%) and were enriched with the CD14^+^CD11b^+^ population. The monocyte-macrophage cells derived from hPBMCs were seeded in 12-well plates at a density of 0.8x10^6^ cells/well. After 24 h of seeding, cells were treated with 10µM of NRP1 inhibitor (EG) at -pre, -during, as well as -post infection phases. Cells were infected with CHIKV at MOI 5 for 2 h. The infected cells were harvested at 8-10 hpi and downstream experiments were conducted. Accordingly, the cells were processed for intracellular staining for CHIKV E2 and supernatants were subjected to ELISA-based cytokine quantification of human TNF. In addition, plaque assay was done from the cell free supernatants to quantify the viral titer.

### 4.14. Western Blot

The cells were harvested 10 hours post-infection and subjected to RIPA lysis. Protease and phosphatase inhibitors were added to the RIPA lysis buffer. The supernatant was collected after centrifuging the cells at 14,000 RPM at 4°C for 40 min. The collected protein lysate was quantified using the BCA assay, and 25 µg of protein was loaded into 10% SDS-PAGE gel wells followed by transfer onto a PVDF membrane. The membrane was blocked using blocking buffer (3% BSA in 1X TBST) before incubation with the primary antibody overnight at 4°C. Primary antibodies against CHIKV E2, CHIKV nsP2, GAPDH, p-p38, p-SAPK/JNK, and NRP1 were used. The corresponding HRP-conjugated secondary antibodies (anti-mouse or anti-rabbit H+L) were used at 1:5000 dilution for 2 h at RT. Blot images were captured using a ChemiDoc XRS+ imaging system. Densitometric analysis was performed using the Image Lab software (Bio-Rad, USA), followed by normalization with respect to housekeeping control i.e., GAPDH. Fold change in band intensities were calculated with respect to mock controls as mentioned previously (5).

### 4.15. Co-immunoprecipitation Assay

For the NRP1-E2/E1 interaction study, RAW 264.7 and HEK cells were infected with CHIKV and lysed with 1X RIPA buffer after viral infection. The lysates were subjected to immunoprecipitation using Dynabeads, as described earlier (5, 52). Briefly, both the mock and CHIKV-infected whole-cell lysates were incubated with pull-down antibody (anti-CHIKV E2 or anti-NRP1) overnight followed by incubation with Dynabeads protein G beads. Post-incubation, the Dynabeads-Ab-Ag complexes were washed, eluted and further processed further for Western blot analysis.

### 4.16. *In silico* molecular docking study

The structural interactions between NRP1 (Human: 2QQM; Mouse: 4GZ9) and CHIKV envelope glycoproteins (E1 and E2; PDB: 3N41) were investigated using a multi-stage *in silico* workflow. Proteins were retrieved via the Schrödinger Get PDB module and standardized using the Protein Preparation Wizard by removing heteroatoms, optimizing H-bonds, assigning pH 7.4 protonation, and refining side chains. Docking was performed using the PIPER algorithm within Maestro. NRP1 served as the receptor, with E1 and E2 defined as ligands. The simulation utilized exhaustive rotational and translational sampling, with final complexes selected based on the lowest PIPER pose energy and the largest cluster size. Binding free energy (ΔG) and dissociation constants (Kd) were predicted using the PRODIGY machine learning framework. The biological relevance of the predicted poses was validated using PRODIGY-CRYSTAL, classifying interactions as either biological interfaces or crystallographic artifacts. Detailed residue-level interactions were analyzed using the Maestro Protein Interaction Analysis module. This included quantifying Interfacial Contacts (ICs) across six residue-type pairs (charged, polar, and apolar) and the calculating link density. Two-dimensional (2D) interaction diagrams and comprehensive interaction tables (CSV) were generated to map specific H-bonds, salt bridges, and hydrophobic contacts at the binding interface. The 3D structures were visualized using UCSF Chimera v1.19, developed by the Resource for Biocomputing, Visualization, and Informatics at the University of California, San Francisco, with support from NIH P41-GM103311 (59).

### 4.17. Antibody Blocking Assay

The anti-NRP1 blocking assay was performed in RAW 264.7 macrophage cells, as per the protocol described previously (5). Cells were incubated with a monoclonal anti-mouse NRP1 antibody or an isotype control anti-rabbit IgG antibody, before incubation with CHIKV at an MOI of 5 for 2 h. Fc receptor blocking solution was used prior to antibody incubation to prevent non-specific IgG binding with macrophages. Cells were harvested at 8hpi and subjected to flow cytometry and western blotting. Cell culture supernatants were analyzed for cytokine response using ELISA. In addition, plaque assay was performed to quantify the viral titer.

### 4.18. siRNA Transfection

The siRNA transfection was performed as per protocol mentioned previously (52). Briefly, HEK293T and C2C12 cells were seeded in 6-well plates and allowed to reach 70% confluency. The C2C12, and HEK cells were transfected with Lipofectamine RNAimax using siRNA corresponding to human (for HEK) or mouse (for C2C12) NRP1 mRNA sequence or with scrambled siRNA as the negative control. No cytotoxic effect was observed in transfected cells. At 24 h post transfection, the cells were infected with CHIKV at MOIs of 0.1 (for HEK), and 0.05 (for C2C12), and harvested at 15hpi (52). The cell lysates were analyzed through western blot and cell-free supernatants were subjected to plaque assay to estimate changes in CHIKV infection.

### 4.19. Plasmid Transfection

Overexpression of NRP1 was performed by using human NRP1 pcDNA3.1 plasmid (Addgene plasmid 213068) as reported earlier (52). The myc-tagged NRP1 (human) was kindly gifted by Yoav Henis Lab, Tel Aviv University, Israel (Addgene plasmid 213068; http://n2t.net/addgene:213068; RRID: Addgene_213068). Briefly, monolayers of HEK293T cells after reaching 70% confluency were transfected with Lipofectamine LTX (Thermo Fisher Scientific, USA) using 1 µg of human NRP1 pcDNA3.1 plasmid or with empty vector as a negative control. No cytotoxic effect was observed in transfected cells. Transfected cells were then infected with CHIKV with an MOI of 0.1. The supernatants and cells were collected after 15 hpi and assayed for plaque assay and Western blot analysis, respectively.

### 4.20. Animal Studies

Animal experiments were conducted in accordance with the guidelines of the Committee for the Purpose of Control and Supervision of Experiments on Animals (CPCSEA) of India with the approval of Institutional Animal Ethics Committee, ILS Bhubaneswar. In brief, 10-12 days old C57BL/6 mice were unbiasedly divided into three groups: mock, infected, and infected-EG treated (n=3 mice per group). The experiment was repeated 3 times. Mice were infected with 1x10^6^ PFU of CHIKV subcutaneously in the flank region of the right hind limb as described previously (33, 60). The control mice (mock) received only the serum free media as mentioned previously (33). The EG-treated group was orally treated with EG (dose: 5 mg/kg body weight) from 2 days before CHIKV infection until 1 day after infection (dpi), at every 24 h intervals. The mock and CHIKV-infected groups received an equal volume of solvent (1X PBS) for the same duration of the study. The dose of EG used in the current study was determined based on previous data showing that a 5mg/kg dose was below the non-toxic threshold and effective in mouse model experiments (27, 54). The mice were monitored for symptoms. At 5 dpi, they were sacrificed followed by harvesting of organs (quadriceps muscle and brain), and collection of serum. The serum cytokines were quantified by cytokine bead array and ELISA based assay. Quadriceps muscle and brain were snap-frozen before lysis in RIPA buffer, followed by homogenisation and sonication to prepare the lysate for Western blot analysis. For plaque assay, the organs were suspended in serum free media followed by homogenisation and filtration with syringe filter (0.22µM). The homogenised solution was centrifuged and the subjected to plaque assay to estimate viral titer.

### 4.21. Data Analysis and Statistical study

The GraphPad Prism 9 software (GraphPad Software Inc., San Diego, USA) was used for statistical analysis. Depending on the experimental design, all comparisons among different groups were performed by either the One-way ANOVA, or the unpaired two-tailed student’s t-test unless otherwise specified. All data were represented as mean ± SD unless stated otherwise. All analyzed data are representative of at least 3 independent experiments with p <0.05 was taken as statistically significant (ns: non-significant, *p <0.05; ** p ≤0.01; ***p ≤0.001; ****p ≤0.0001).

## DATA AVAILABILITY

The data that support the findings of this study have been included in the manuscript as main figures or supplementary files.

## ACKNOWLEDGEMENTS

We are thankful to the NISER Central Instrumentation and Flow Cytometry Facility for providing pivotal support to conduct this study. We thank the health centre of NISER, Bhubaneswar, for helping with the human PBMC experiments and Animal house facility of Institute of Life Sciences (ILS) for the *in vivo* experiments. We acknowledge Dr. Abdur Rahaman’s lab, SBS, NISER, for their help with the overexpression work. We thank Rajshree Rajmohan Jena for his help with *in vivo* experiments. We acknowledge Hemendra M, Ayan Chakraborty, Shoumyodeep Barman, Parthasarathi Jena, Supriya Suman Keshry, and Agniv Chakraborty for their help with the experiments and valuable suggestions to improve the manuscript.

## AUTHOR CONTRIBUTIONS (CRediT)

**KST:** Conceptualization, Investigation, Formal Analysis, Methodology, Validation, Writing-Original Draft, Writing-Reviewing & editing. **CM:** Methodology, Investigation, Formal Analysis, Writing-Reviewing & editing. **SG:** Methodology, Investigation, Formal Analysis, Writing-Reviewing & editing. **KM:** Methodology, Writing-Reviewing & editing. **SS:** Methodology, Investigation. **BB:** Methodology. **SK:** Writing-Reviewing & editing. **HB:** Methodology. **MG:** *in silico* Methodology. **BBB:** *in silico* Methodology, Investigation, Validation, Writing-Reviewing & editing. **SoC:** Resources, Supervision, Writing-Reviewing & editing. **SuC:** Funding Acquisition, Project administration, Conceptualization, Investigation, Resources, Supervision, Writing-Reviewing & editing.

